# Assessing Cognitive Functions Remotely Using a Music-Game-Like Program

**DOI:** 10.1101/2024.04.07.588461

**Authors:** Maxime Perron, Ashna Imran, Rebecca Alain, Claude Alain, Yayoi Sakaki

## Abstract

There is a growing need to develop ways of assessing cognitive functions remotely and providing interventions using a web-based approach. The Ipsilon Test is a music-based cognitive assessment and training tablet application. On each trial, it presents simplified musical notation with colours, spatial cues, and gestural references so that users can translate spatial information into motor actions by tapping on the screen. In this study, we examined the correlation between the performance of a group of healthy older adults on the Ipsilon test and standard measures of cognitive function—a total of 30 participants aged 58 and over were recruited for the study. Before and after one week of training using the web-based program Ipsilon, all participants completed an online version of the Montreal Cognitive Assessment (MoCA) and the visual Stroop task. Of the participants recruited, 22 participants completed the Ipsilon training and cognitive testing. In our sample, performance on the Ipsilon test was generally high, with all participants scoring above the chance level. Importantly, we found a correlation between Ipsilon test performance and pre-Ipsilon MoCA scores. In addition, participants who scored higher on the Ipsilon test also showed improvement in the visual Stroop task after Ipsilon training, particularly in the ability to inhibit irrelevant information. These results suggest the Ipsilon test is a practical web-based cognitive assessment and training tool. Future research will explore its relationship with other cognitive tests and its diagnostic power for differentiating individuals with normal cognition from those with cognitive disorders.

## Introduction

The urgent need to transition non-acute care to remotely operable solutions due to the COVID-19 pandemic and the necessary risk mitigation have fueled the demand for digitized solutions. Over the last few decades, an increasing number of studies have been conducted to determine the feasibility of computerized medical devices and software in cognitive care, especially in the assessment and training domains (Björngrim et al., 2019; Dahmen et al., 2017; Kessels, 2019; Vermeent et al., 2020). While additional validations remain necessary, findings from these studies indicate that digitized solutions are easy to use and provide comparable diagnostic performance to the original paper-pencil versions.

Recent studies have shown that lifelong engagement in musical activities may help slow cognitive decline (Hanna-Pladdy & Gajewski, 2012; Parbery-Clark et al., 2011; Tremblay & Perron, 2023; Zendel & Alain, 2012), maintain brain health closer to a younger state (Rogenmoser et al., 2018), and preserve or even enhance neural functioning in old age (Alain et al., 2014; Bidelman & Alain, 2015; Moussard et al., 2016). For example, older musicians perform better than non-musicians in several perceptual and cognitive tasks, including speech in noise perception (Zendel & Alain, 2012), concurrent sound segregation (Zendel & Alain, 2013), phonemic fluency, verbal working memory, immediate recall, visuospatial judgment, motor dexterity (Hanna-Pladdy & Gajewski, 2012), and inhibitory control (Moussard et al., 2016; Tremblay & Perron, 2023). Evidence also suggests that short-term musical training (e.g., piano lessons) can improve working memory and other executive functions (Alain et al., 2019; Bugos et al., 2022). Altogether, this indicates that engaging in musical activities at any stage of life can have significant cognitive benefits.

This study aims to evaluate the Ipsilon Test (IPS-001) as a practical telehealth cognitive care solution for assessing cognitive impairment and providing rehabilitation. The Ipsilon Test is a gamified cognitive functionality training and assessment application using musical playing to generate the responses for analysis. It is a remotely operated self-care tool that is easy to use on a tablet device (e.g., iPad). The Ipsilon Test collects the user’s time-stamped finger-tapping response, and the number of accurately answered responses. It exhibits simplified music notation with colour, spatial, and gestural references for the users to translate spatial and motor (e.g., which hand to use with the corresponding areas on the piano grid) information into motor actions and deliver a response by simply tapping the screen. The language- and auditory-free nature of the Ipsilon Test could make it a versatile cognitive assessment tool suitable for measuring cognition in various populations.

To establish the Ipsilon Test proof of concept, we asked a group of older adults to perform the Ipsilon training. Then, we examined how the performance on the Ipsilon test correlated with standard measures of cognitive functions. Specifically, we correlated Ipsilon test performance with the MoCA score, a widely used assessment tool for evaluating general cognitive functioning and diagnosing mild cognitive impairment. We also investigated the relationship between the Ipsilon Test and executive function using the Stroop task, which measures the ability to inhibit automatic responses. We expected to observe correlations between performance on the Ipsilon Test and these cognitive measures, which would provide initial evidence supporting using the Ipsilon Test as a cognitive assessment tool.

## Method

### Participants

Seventy-four individuals filled out the “Consent Form” after being recruited from Baycrest’s participant database, Honeybee Hub (https://hello.honeybeehub.io/), and word of mouth. Eligibility was verified through online questionnaires on Qualtrics (Qualtrics, Provo, UT), and 30 older adults were eligible to participate. Participants have been screened for a history of neurological or psychiatric disorders, medications that hinder cognitive abilities, functional independence, colour blindness, uncorrected vision, and hearing loss. Of this sample, a total of 23 participants completed the Ipsilon Test. Of the 23 participants, 19 completed all sessions and cognitive measures. Three experienced technical issues with the computerized tests, and one chose not to continue with the study. Only participants with at least one pre- and post-Ipsilon cognitive measure were included in the final sample (*N* = 22). Participants ranged in age from 58 to 86 years (*M* = 71.3, *SD* = 8.4) and included 17 females and five males. The Baycrest Research Ethics Board approved the project, and participants were compensated for their time.

### Procedure

Participants were asked to take the Ipsilon Test and to complete pre- and post-Ipsilon cognitive assessment. Participants were asked to devote 15 minutes daily to the Ipsilon Test for at least five days to ensure regular engagement. The cognitive assessment included the Montreal Cognitive Assessment (MoCA) (Nasreddine et al., 2005) and a computerized version of the classic visual Stroop task (Stroop, 1935). The MoCA was administered by Zoom or telephone, depending on the participant’s chosen mode of communication. Trained research assistants administered the MoCA assessment remotely. For participants using Zoom, the classic version of the MoCA was completed, with scores up to 30 points. Participants who opted for telephone administration completed the telephone version of the MoCA with scores of up to 22 points. To facilitate comparisons, scores were standardized by calculating the percentage ratio. The visual Stroop task was performed using participants’ own devices on Inquisit software (Millisecond Software, 2020). Participants received assistance throughout the process to ensure they understood and complied with all the steps.

### Ipsilon Test

The participants were provided with a de-identified, study-specific email addresses and password for registration to the Ipsilon Test website. After completing the registration process, they were given a brief tutorial on how to use the application. The test consists of nine levels, each consisting of ten trials. The levels are divided into odd and even numbers, where odd-numbered levels involve right-hand use and even-numbered levels involve left-hand use. The difficulty of the spatial identification task increases as the level number increases, with Levels 1 and 2 being the simplest. Level 9 is unique in that it selects random patterns from Levels 1 through 8. Before starting each data collection session, participants were given a practice run of ten trials to familiarize themselves with the task. During each trial participants were presented with two black dots on a white background, similar to how the music notation works but simpler, accompanied by colored lines representing piano keys underneath (Figure 1). The spatial-motor instructions were displayed using these visual cues. Participants were then required to reproduce the notes represented by the two black dots in the same order by clicking on the corresponding notes on a ten-note piano keyboard displayed below. Once the participant answered, a three-second visual countdown would commence before the next trial began. Participants were only allowed to progress to the next level if they achieved a minimum accuracy of 80%. Performance on the Ipsilon Test was assessed based on three measures: Accuracy (percentage of error), its variability (standard deviation), mean reaction time (measured in milliseconds), and its variability (SD).

**Figure 1.**
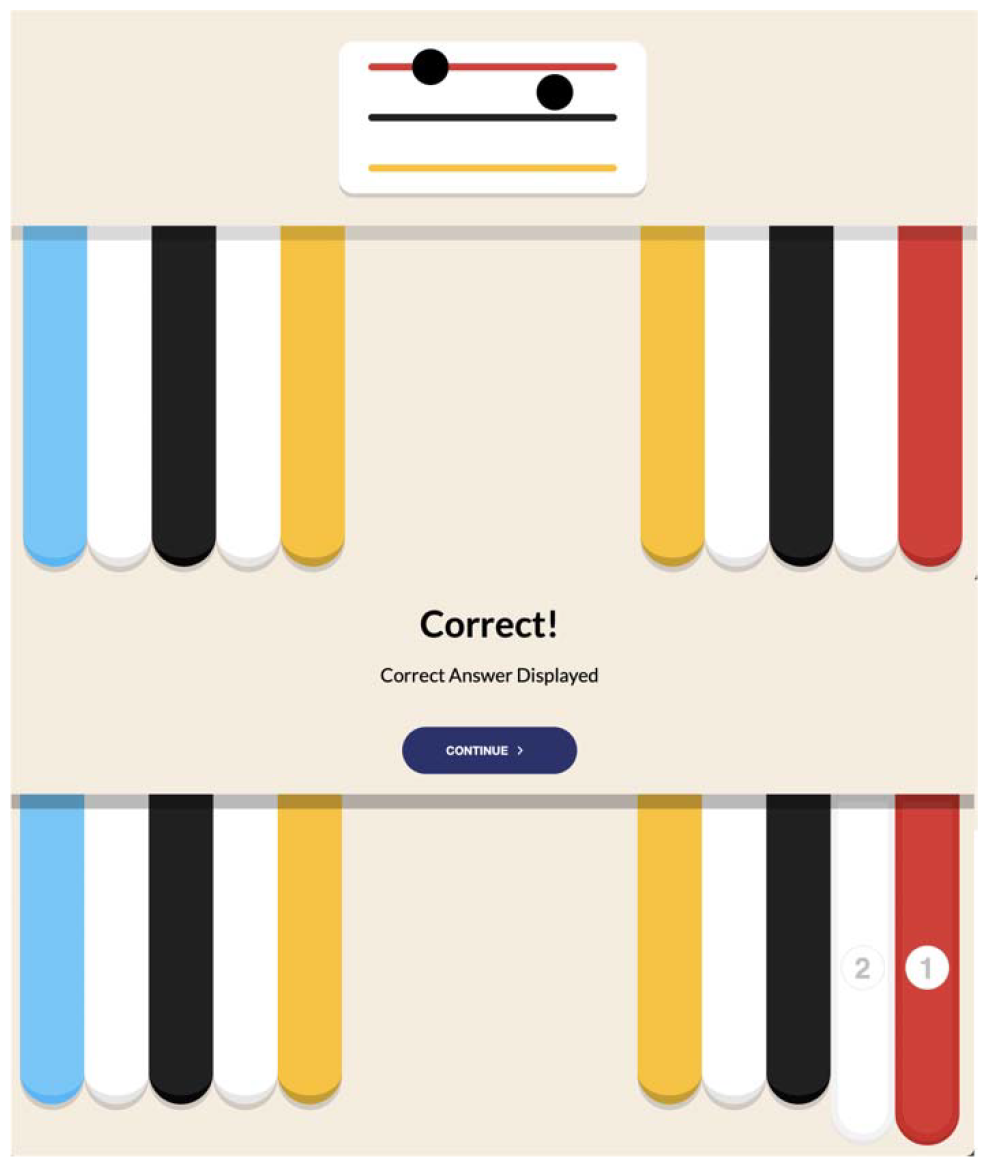
Example of a practice trial. The top image shows sample visual stimuli as spatial-motor instructions and colour-referenced piano grids for response collection. The bottom image shows feedback in the event of a correct response.

### Visual Stroop task

Before and after completing all levels of the Ipsilon Test, participants completed the visual version of the Stroop task (Stroop, 1935). The task was implemented by Millisecond Software and then modified by our team for the current study. Participants were asked to press a button indicating the colour in which the words “red,” “green,” “black,” or “blue” were printed as quickly and accurately as possible. On any given trial, the word “red,” “green,” “black,” or “blue” printed in red, green, black, or blue characters was presented against a white background. The task comprised three types of trials: congruent, incongruent, and neutral. In congruent trials, the printed word and the colour in which it was printed matched (e.g., the word “blue” printed in blue). In incongruent trials, the printed word and the printed colour did not match (e.g., the word “blue” was printed in black). In neutral trials, a rectangular shape printed in red, green, black, or blue was presented, and participants were asked to indicate the shape’s colour. Performance in the Stroop task was measured using the interference and facilitation effects. The interference effect measures the delay in reaction time when the colour of a word is incongruent with the meaning it represents. It is calculated by subtracting the reaction time of the neutral condition from that of the incongruent condition (Interference = Incongruent - Neutral). The higher the scores, the greater the interference (i.e., the poorer the inhibitory control). Conversely, the facilitation effect translates into improved response time when word and colour are congruent. The facilitation effect is calculated by subtracting the RT of the neutral condition from that of the congruent condition (Facilitation = Congruent - Neutral). The lower the scores, the greater the facilitation (i.e., better inhibitory control).

### Statistical analyses

Data were analyzed using R Studio version 4.1.1 (RStudio Team, 2022). As most variables were not normally distributed, non-parametric statistics were used. First, we performed descriptive statistics for performance within the Ipsilon Test (accuracy, mean reaction time, reaction time variability). Next, we used paired Wilcoxon signed-ranks tests to compare performance on the MoCA and the Stroop visual task before and after Ipsilon training using the wilcox_test function in the rstatix package version 0.7.2. Finally, we performed Spearman correlations between Ipsilon performance and cognitive measures before and after Ipsilon training using the rcorr function in Hmisc package version 5.1.0. An alpha level of 0.05 was used for all analyses to determine the statistical significance.

## Results

### Overall performance within the Ipsilon test

The participants exhibited an average accuracy rate of 84% (*SD* = 12), indicating a relatively high level of task performance. All participants demonstrated above-average performance, with accuracy ranging from 66% to 97%. Reaction times varied between 2010 and 6014 ms, with an average of 3832 ms (*SD* = 1024). The variability in reaction times ranged from 449 to 2703 ms, with an average of 1339 ms (*SD* = 686).

### Difference in cognitive performance before and after Ipsilon Test

Wilcoxon signed-ranks tests were conducted to compare the MoCA and Stroop task performance before and after the Ipsilon Test. The results revealed a significant decrease in interference effects after the Ipsilon Test (*V* = 120, *p* = 0.04). However, no significant differences between the two sessions were observed in the MoCA score (*V* = 51.5, *p* = 0.98) and facilitation Stroop effect (*V* = 134, *p* = 0.12).

### Correlation between cognitive performance and Ipsilon performance

Spearman correlations were conducted to examine the relationships between Ipsilon performance metrics and pre- and post-Ipsilon performance on the MoCA and visual Stroop task. The results are shown in Figure 2. First, a moderate negative correlation was observed between the pre-Ipsilon MoCA score, mean reaction time (*r* = -0.48, *p* = 0.02) and reaction time variability (r = -0.46, *p* = 0.03). These results suggest that individuals with lower MoCA scores prior to the Ipsilon Test tend to exhibit longer reaction times and greater variability in their reaction times during the Ipsilon Test. Furthermore, the post-Ipsilon interference effect negatively correlated with Ipsilon Test accuracy (r = -0.49, *p* = 0.03). This indicates that individuals who demonstrate higher accuracy during the Ipsilon Test also exhibit better inhibitory control after completing the Ipsilon Test.

**Figure 2.**
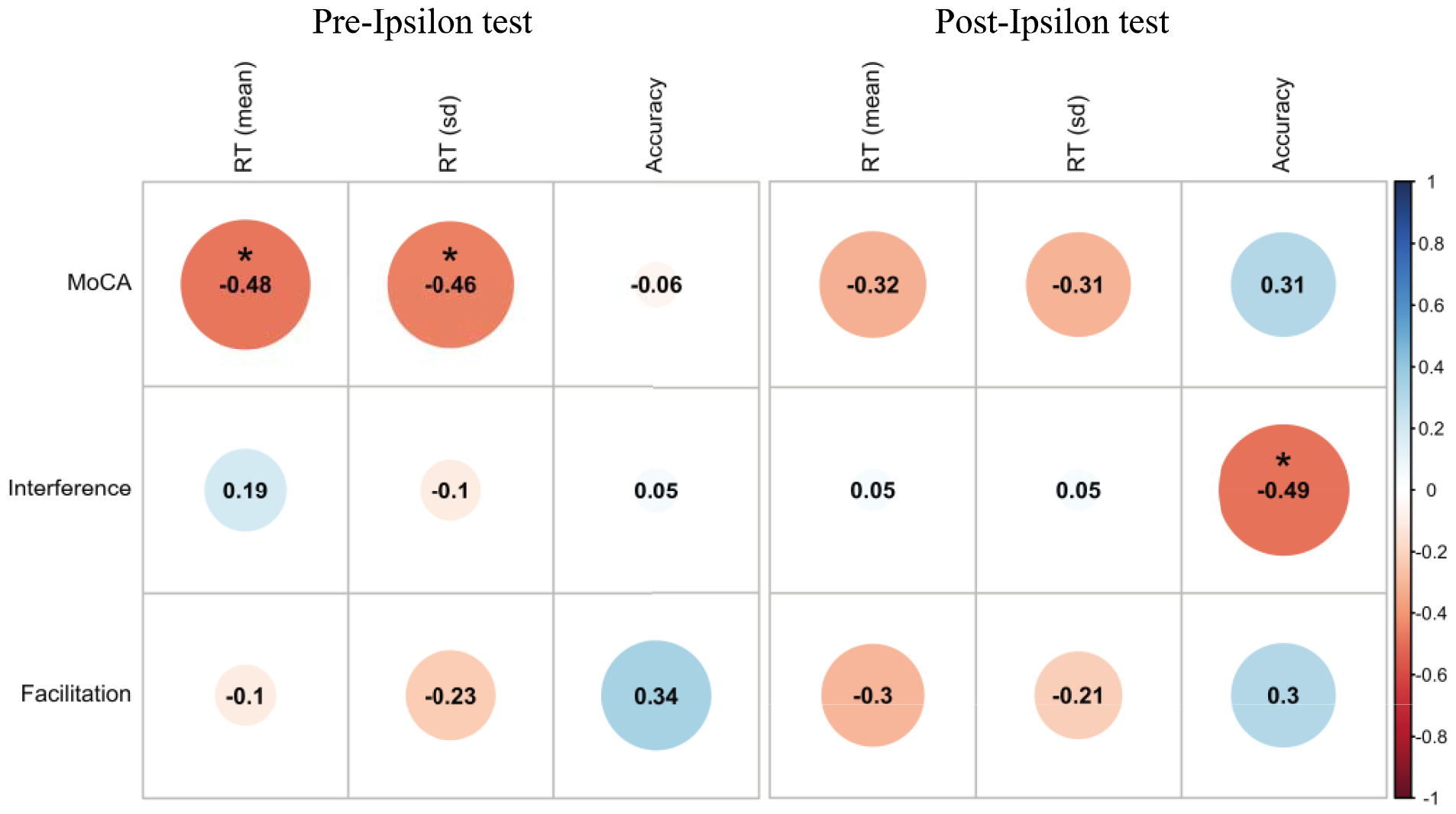
Spearman correlations between Ipsilon performance metrics (accuracy, mean reaction time, reaction time variability) and pre- and post-Ipsilon performance on the MoCA and visual Stroop task.

## Discussion

The Ipsilon Test has been developed to provide a web-based cognitive assessment and training program. It is based on a simulated piano game that enables individuals to train their cognition while assessing the possibility of developing cognitive decline. As a first step towards this goal, we examined, from a group of cognitively healthy older adults, how performance at the Ipsilon Test related to other standard measures of cognitive function. Specifically, we focused on two cognitive measures: the MoCA and the Stroop task. We used the MoCA to determine whether Ipsilon test performance could indicate general cognitive functioning. The MoCA is widely used to assess global cognitive function and diagnose mild cognitive impairment. We also used the Stroop task as a more direct measure of executive function, which corresponds closely to the cognitive abilities assessed by the Ipsilon Test. The Stroop task is designed to assess executive functions, mainly inhibitory control.

Our findings revealed interesting correlations between the MoCA scores and Ipsilon test performance. We observed a moderate negative correlation between pre-Ipsilon MoCA scores and mean reaction time, as well as reaction time variability during the test. This indicates that people with lower MoCA scores before taking the Ipsilon Test tend to exhibit slower and more inconsistent reaction times during the test. Thus, the MoCA score appears to have predictive value for Ipsilon test performance, with lower scores indicating slower and more variable reaction times. Furthermore, we observed a significant reduction in the Stroop interference effect after the Ipsilon training. The Stroop interference effect results from difficulties in suppressing irrelevant information, and the observed reduction suggests that Ipsilon training positively influenced participants’ inhibitory control abilities. Specifically, participants demonstrated an improved ability to ignore distractions and focus on task-relevant stimuli following the Ipsilon training. In addition, we found a negative correlation between the post-Ipsilon interference effect and the Ipsilon Test accuracy. This suggests that individuals who demonstrated greater accuracy during the Ipsilon Test also showed better inhibitory control after the training was completed. Therefore, good performance on the Ipsilon Test may indicate improved inhibitory control abilities, resulting in reduced interference effects.

However, it is important to recognize the limitations of our study. The absence of a control group makes it difficult to definitively attribute the observed reduction in the interference effect to the Ipsilon training, as opposed to the effects of repeated exposure to the task. Nevertheless, the lack of correlation between pre-Ipsilon interference effects and Ipsilon test performance suggests that this reduction may be an effect of the Ipsilon training. Another limitation of our study is the small sample size, which may have impacted statistical power. However, it is encouraging that we found moderate correlations between Ipsilon performance and standard cognitive measures.

## Conclusion

In conclusion, our study demonstrates the feasibility of using the Ipsilon Test as a cognitive training tool in cognitively healthy older adults. Nevertheless, further studies involving larger and more diverse samples are needed to confirm its diagnostic accuracy. In addition, further research is needed to determine its suitability for diverse populations, such as individuals with mild cognitive impairment or Alzheimer’s disease. The Ipsilon test has the potential to become a valuable option for online cognitive assessment and training.

